# LIPUS attenuates knee joint capsule fibrosis and athrogenic contracture through the TGF-β1/Smad signaling pathway

**DOI:** 10.1101/2023.03.16.532928

**Authors:** Ting Zhou, Feng Wang, Yun Zhou, Chen Xu Zhou, Quan Bing Zhang

## Abstract

As one of main causes of athrogenic contracture, joint capsule fibrosis which is described as a condition with excessive deposition of collagen components and extracellular matrix (ECM) in joint capsule, is a response to long-time immobilization. The purpose of this study was to explore the effect and the underlying mechanism of low-intensity pulsed ultrasound (LIPUS) in treating knee joint capsule fibrosis. A rabbit model of knee joint capsule fibrosis induced by 6w-immobilization was employed in this study. The degree of knee joint capsule fibrosis was assessed by measurement of arthrogenic contracture and Masson-staining. Furthermore, malondialdehyde (MDA) and superoxide dismutase (SOD) were measured to assess the level of reactive oxygen species (ROS). Apart from these, the activation of TGF-β1/Smad signaling pathway was determined through western blot analysis contained TGF-β1, Smad2, p-Smad2, Smad3, p-Smad3 and Smad4, and immunohistochemical staining for p-Smad2/3 positive cells. After 6 wk-immobilization, the degree of arthrogenic contracture and the collagen density were increased. Moreover, the activity of MDA was upregulated and the content of SOD was downregulated. Correspondingly, the TGF-β1/Smad signaling pathway was significantly activated. After 2 wk-LIPUS treatment, the degree of arthrogenic contracture and the collagen density were lower than 2 wk-remobilizaiton. Relatively, the activity of MDA was decresed and the content of SOD was increased compared with 2 wk-remobilizaiton. Importantly,the TGF-β1/Smad signaling pathway was significantly inhibited compared with 2 wk-remobilizaiton. Our findings thus conclude that high level ROS and hyperactive TGF-β1/Smad signaling pathway might be one of the causes of knee joint capsule fibrosis induced by immobilization and LIPUS attenuated the severity of immobilization-induced knee joint capsule fibrosis through inhibition of the production of ROS and the activation of TGF-β1/Smad signaling pathway. Our findings might provide a theoretical basis for knee joint capsule fibrosis after immobilization and provide the potential therapeutic target.

## 1. Introduction

Knee joint contracture is a condition associated with a decline in range of motion (ROM), which limits the patient’s activities of daily living (ADL) by impairing movement ability and joint function (Chen et al. 2022; Yang et al. 2021). While advances in understanding of disease mechanisms have improved in recent years, knee joint contracture remains remarkably extremely difficult to treat. Myogenic and arthrogenic contracture are the two structural components contributing to joint contracture formation and it is important that the joint capsule is regarded as the critical motion-limiting anatomic structure in a developing arthrogenic contracture (Wang et al. 2019). Furthermore, during remobilization, although myogenic contracture recovers spontaneously, arthrogenic contracture is irreversible or deteriorates further (Kaneguchi et al. 2018). To prevent the progression of arthrogenic contracture during remobilization, therefore, effective therapy should be taken to control joint capsule fibrosis.

Joint capsule specimens from contracted joints are thicker and less compliant than those from unaffected joints (Pujol et al. 2015). The condition of joint capsule fibrosis, characterized by excessive collagen deposition and extracellular matrix (ECM), is the result of prolonged immobilization of joints (Kaneguchi et al. 2018). Our previous study revealed that transforming growth factor-beta1 (TGF-β1) was an important regulator of joint capsule fibrosis. As time progressed in a rabbit model of knee contracture in extension, arthrogenic contracture developed, possibly because of fibrosis in the joint capsule (Zhou et al. 2020). Recently, some studies indicated that the activation of TGF-β1/Smad signaling plays an important role in arthrogenic contracture and joint capsule fibrosis (Xiao et al. 2020). TGF-β1 has a substantial influence in the process of fibrosis development characteristic of joint capsule, which produced by fibroblast promotes fibroblast differentiation into myofibroblasts and produces a large amount of ECM components (Mao et al. 2021). Therefore, one of the major factors during joint capsule fibrosis development is TGF-1 expression. Smad2/3 was one of the major downstream targets mediated the fibrogenic activity of TGF-β1. In fibroblasts, phosphorylation of Smad2/3 mediates TGF-β1 regulation of myofibroblast differentiation. Smad2/3 is phosphorylated and interacted with Smad4 to form a complex that enters the nucleus and regulates -SMA, Collagen I, and Collagen III gene expression (Wang et al. 2022).

Notably, joint immobilization has been shown to induce hypoxic conditions in knee joint (Huang et al. 2021; Yabe et al. 2013). Under hypoxic conditions, reactive oxygen species (ROS) increase significantly in joint capsule (Jiang et al. 2018). Besides being crucial to the survival of cells and normal communication between cells, ROS can also be causative of fibrosis. It has been found that ROS also contribute to myofibroblast differentiation, mediating TGF-1 signaling pathways and interacting with extracellular matrix (Siani et al. 2014). Previous research demonstrated that high ROS generation not only elevated the expression of TGF-β1 but also increased the fibrosis protein markers including collagen I and α-SMA in synovial myofibroblasts after knee joint immobilization (Jiang et al. 2018). Thus, in order to improve arthrogenic contracture and joint capsule fibrosis should include inhibition of the production of ROS and the activation of TGF-1/Smad signaling pathway in joint capsule.

Low-intensity pulsed ultrasound (LIPUS) is a type of ultrasound that emits pulsed waves at lower intensities (Xin et al. 2016). Considerable previous experiments demonstrated that LIPUS acted as a role of ROS scavenger. Recent studies have found that LIPUS treatment reduces ROS, apoptosis, and matrix metalloproteinases (MMPs), thus alleviating articular cartilage damage in knee osteoarthritis (Jiang et al. 2019). Furthermore, LIPUS prevented endothelial-mesenchymal transitions by decreasing ROS accumulation and enhancing PI3K signaling cascade activation in human aortic endothelial cells. Importantly, the role of LIPUS in anti-fibrosis is well documented in myocardial infarction, osteoarthritis and hypertensive nephropathy, etc (Liao et al. 2021; Zuo et al. 2021; Zhao et al. 2021). However, there are currently no previous reports investigating the curative effect of LIPUS on arthrogenic contracture and joint capsule fibrosis during immobilization or remobilization. Given the established functions of LIPUS in previous studies, we hypothesized that LIPUS treatment effectively attenuates the progression of joint capsule fibrosis via inhibition of ROS mediated TGF-β1/Smad signaling pathway during knee joint remobilization (Figure 1. A). The current study was, therefore, designed to preliminarily investigate the effect of LIPUS on arthrogenic contracture and joint capsule fibrosis in our established rabbit model of extending knee joint contracture (Zhou et al. 2020), and to investigate the possible mechanism of action. This may provide a potential theoretical basis and experimental support for LIPUS in the treatment of arthrogenic contracture and joint capsule fibrosis.

**Figure 1.**
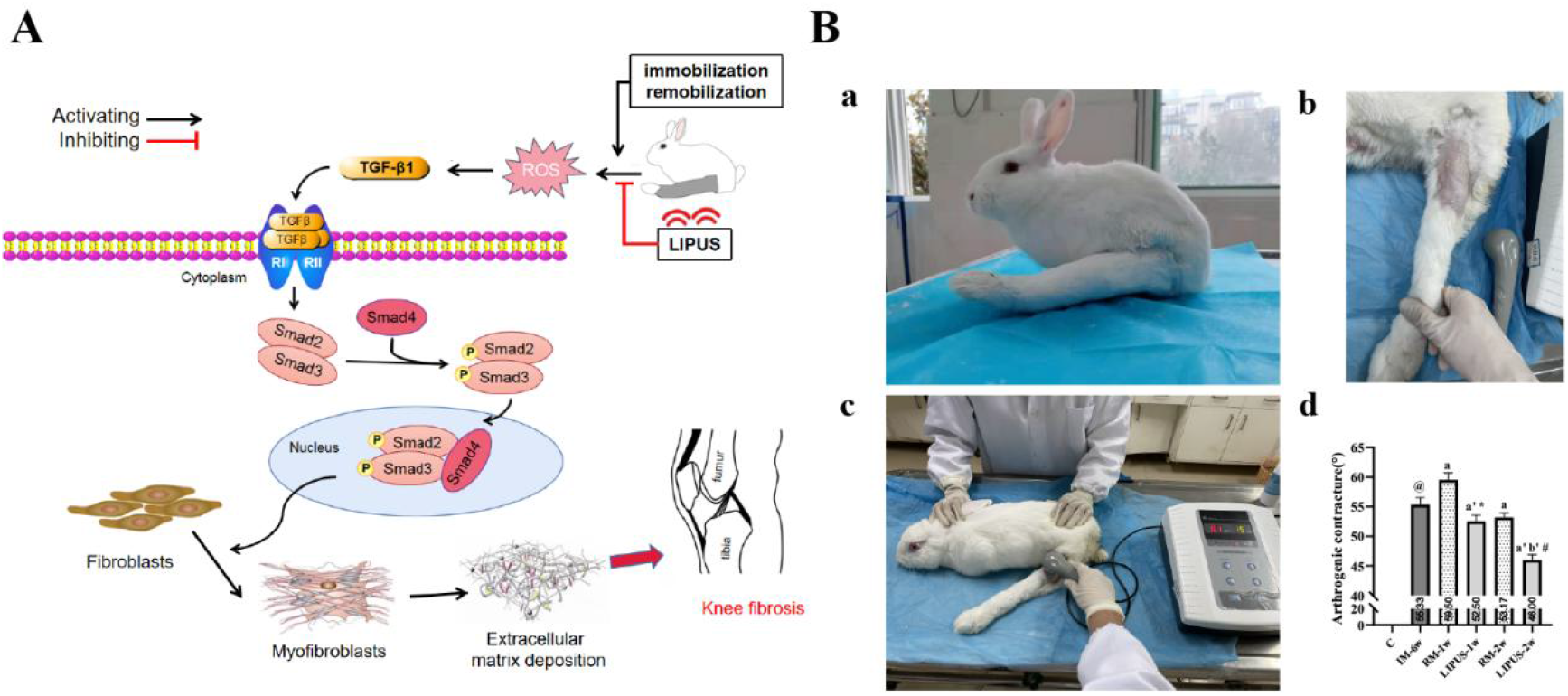
(A) Research hypothesis. (B) Animal model and intervention methods. (a) Unilateral immobilization of the rabbit knee joint at full extension using a plaster cast from the groin to the proximal toes. (b) The left knee joint after 6 wks-immobilization. (c) The handpiece of low-intensity pulsed ultrasound device and the rabbit left knee joint coated with couplant. (d) Effects of remobilization and LIPUS on the arthrogenic contracture (°) in each group (n = 6). ^@^ *P* < .05 vs. group C, ^a^ *P* < .05 vs. group IM-6w, ^b^ *P* < .05 vs. group RM-1w, ^a’^ *P* < .05 vs. group IM-6w, ^b’^ *P* < .05 vs. group LIPUS-1w, ^*^ *P* < .05 vs. group RM-1w, ^#^ *P* < .05 vs. group RM-2w.

## 2. Methods

### 2.1 Grouping and Animal model

Animal care and experimental procedures were performed in accordance with the Guidelines for Animal Experimentation of Anhui Medical University and were approved by the Institutional Animal Care and Use Committee (LLSC20190761). Thirty-six male New Zealand white rabbits (approximately 4-month-old, 2000g-2500g) were provided from Anhui Medical University Experimental Animal Center (Hefei, China) and randomly assigned to control (group C, rabbits without any intervention, n = 6), immobilization (group IM-6w, for six-week immobilization, n = 6 each), remobilization (group RM-1w and group RM-2w, for one-week and two-week remobilization after six-week immobilization, respectively; n = 6 each), and LIPUS treatment (group LIPUS-1w and group LIPUS-2w, for one-week and two-week remobilization combined LIPUS treatment after six-week immobilization, respectively; n = 6 each) groups.

According to our previous study (Huang et al. 2021; Zhou et al. 2020; Zhang et al. 2018), the rabbit left knee joint was fixed for 6 weeks at full extension using a plaster cast from groin to proximal toes to established arthrogenic contracture model (Figure 1. B. a). All rabbits were reared in the same environment and housed 1 per cage. Specifically, they were housed one per cage at 22°C-24°C with a 12-hr light/dark cycle and were allowed access to food and water ad libitum. In each group, during the experimental period, the area other than the immobilized part was left free to move inside the cage.

### 2.2 LIPUS treatment

Rabbits were treated using a LIPUS device (HS-501, HANIL-TM, Korea) under a frequency of 1 MHz at an intensity of 0.1W/cm^2^ (Figure 1. B. c) with a pulse duration of 15s. To avoid ultrasound energy attrition, the left knee was shaved to remove all fur surrounding the knee joint, and the treatment area was selected both the inside and outside of the left knee joint (Figure 1. B. b). Ultrasound coupling gel was applied to the knee to insure acoustic transmission. Each treatment lasted 15 minutes, once a day until the end of the experiment. During this time the treatment probe was gently moved across the treatment area to prevent skin damage or irritation from prolonged treatment at a single location. Afterwards the ultrasound gel was wiped from the knee, and the rabbit was returned to its cage. We occasionally observed signs of distress among the rabbits during our treatment, but every time this occurred, it appeared to be due to the rabbit resisting the restraint rather than reacting to the treatment itself. It was possible to pause the treatment until the rabbit was calmed as needed. These breaks lasted no longer than a minute.

### 2.3 Assessment of arthrogenic contracture

At end point, the rabbits were killed with an overdose of sodium pentobarbital. The left hindlimb was dislocated at the left hip joint and completely removed. As in our previous study, arthrogenic contracture = ROM after myotomy (control knee) - ROM after myotomy (experimental knee), the range of motion (ROM) after knee myotomy was measured in the control group and the experimental group was measured, respectively. In accordance with the method described by our previous study (Zhou et al. 2020; Zhang et al. 2018), we evaluated the arthrogenic contracture caused by the articular structures.

### 2.4 Anterior knee joint capsule preparation

Research on the effect of joint immobilization has identified the anterior knee capsule as a key structure limiting knee ROM in flexion (Zhou et al. 2020; Zhang et al. 2018). A reduction in anterior knee capsule length was positively correlated with a decrease in flexion. After the measurements of ROM, the anterior joint capsule was harvested from the knee and partitioned into equal samples. One part of samples were immediately detected for MDA and SOD Levels, and other samples were snap frozen in liquid nitrogen and then stored at -80 °C.

### 2.5 Determination of MDA and SOD Levels

ROS levels are balanced by ROS-generating enzyme, NADPH oxidase, and antioxidant enzymes such as superoxide dismutase (SOD), catalase, and glutathione peroxidase (GPx) (Moloney et al. 2018). ROS can activate antioxidant enzymes and excessive ROS can react with membrane lipids to generate malondialdehyde (MDA) (Qiu et al. 2019). Therefore, SOD and MDA is commonly used for evaluating the level of ROS. As previous research (Ding et, al. 2022), the anterior knee joint capsule was added with nine times the volume of normal saline with the proportion of weight (g): Volume (ML) = 1:9 to prepare the 10% tissue homogenate at 2500 rpm for 10 min using a commercial kit (Nanjing Jiancheng Bioengineering Institute, China). The levels of malondialdehyde (MDA, A003-1) and superoxide dismutase (SOD, A001-3) in the tissue homogenate were determined according to the manufacturer ’ s protocol.

### 2.6 Masson Staining

Masson staining was used to evaluate joint capsule fibrosis with a commercially available kit (Solarbio, China). The samples of anterior knee joint capsule sections were baked at 60°C for 90 min, immersed respectively in xylene (3 min × 3 times), absolute ethanol (3 min × 2 times), 95% and 75% ethanol (each for 3 min), and flushed by running water. Then the sections were stained by hematoxylin for 10 min, flushed by running water, differentiated by hydrochloric acid for several seconds, flushed by running water again and colorized to blue by ammonia for several minutes. Next, ponceau-acid fuchsin, 12 molybdophosphoric acid solution and green staining solution were successively used to stain the sections for 4 ∼ 20 s, 2 ∼ 4 min, and 2 ∼ 5 min, respectively. After washed by water and baked in an oven, the sections were sealed by neutral gum, observed and photographed under the microscope. In each section, 6 fields of vision were chosen for observation under the microscope (× 100). Image J was employed to perform semi-quantitative analysis: the percentage of the area of collagenous fibrosis (presenting in blue color) in the fields of vision accounting for the total area of the field was measured and the average value was taken. Three independent investigators (experienced pathologists) confirmed the quantification of each slide.

### 2.7 Immunohistochemistry

Joint capsule specimens were cut to 3–5 μm-thick sections. The sections were dewaxed and rehydrated in mixture of xylene and ethanol. Then, the sections were incubated with anti-phospho-smad2/3 (Wanleibio, WL02305, dilution, 1:150) at 4°C over night. The sections were treated with PBS solution before being incubated with peroxidase-conjugated goat anti-rabbit IgG-HRP (Abcam, ab205718, dilution, 1:1000) for 20 minutes at 37°C. A diaminobenzidine (DAB) colorimetric method was used to assess the activity. Then hematoxylin was used to counterstain the sections. Under the microscope, 6 fields of vision were examined in each section (× 400). From each set of six randomly analyzed fields, an average number of positive cells was calculated using Image J. Three independent investigators (experienced pathologists) confirmed the quantification of each slide.

### 2.8 Western blot

The joint capsule samples were ground into powder with liquid nitrogen using a grinder and homogenized in RIPA buffer (Beyotime, China) containing protease and phosphatase inhibitor cocktail (Beyotime, China) at 4 °C. Homogenates were centrifuged at 12,000 × g for 30 min three times at 4 °C, and the resulting supernatants were collected. By using the bicinchoninic acid method, protein concentrations were determined. On a 10% sodium dodecyl sulfate-polyacrylamide electrophoresis gel, protein lysates were separated and transferred onto polyvinylidene fluoride membranes (Millipore, USA). After being blocked with five-percent non-fat dry milk in Tris-buffered saline Tween-20 (TBST) at room temperature for two hours, the membranes were respectively incubated with GAPDH (Affinity, AF7021, dilution, 1:1000), rabbit anti-TGF-β1 (Bioss, bs0103R, dilution, 1:500), rabbit anti-Smad2 (Bioss, bs-0718R, dilution 1:1000), rabbit anti-phospho-Smad2 (Bioss, bs-24530R, dilution, 1:1000), rabbit anti-Smad3 (1:1000, Bioss,bs-3484R), rabbit anti-phospho-Smad3(Bioss, bs-19452R, dilution,1:1000), rabbit anti-Smad4(1:1000, Bioss, bsm-52225R), rabbit anti-CollagenI (Bioss,bsm-52225R, dilution, 1:1000), rabbit anti-CollagenIII (Bioss, bs-0549R, dilution, 1:1000), rabbit anti-α-SMA(Bioss,bs-10196R, dilution, 1:500) at four degrees Celsius overnight. On the second day, after being washed in TBST solution three times for 10 min per wash, the membranes were incubated with peroxidaseconjugated affinipure goat anti-rabbit IgG-HRP (Abcam, ab205718, dilution, 1:5000) as the secondary antibody for two hours at room temperature.

Following three TBST washes for 10 minutes per wash, the membranes were evaluated with the enhanced chemiluminescence system, as directed by the manufacturer. The band densities were quantified using Image J software.

### 2.9 Statistical analysis

All data are expressed as the mean ± standard error of the mean. The assumptions of normality of data and homogeneity of variances between the groups were analyzed by SPSS 25.0 (IBM Corp, Armonk, NY, USA). Unpaired t-tests were performed to compare group C and group IM. The effect of intervention (LIPUS and remobilization) for all outcome measures were evaluated by two-way repeated measures analysis of variance (ANOVA) with group as a between-subjects factor and time as a within-subject factor. A simple main effect test after Bonferroni correction was performed as a post hoc analysis when the interaction effect was significant. All statistical graphs were processed using GraphPad Prism 8 (GraphPad Scientific, San Diego, CA, USA).

## 3. Results

Repeated measures ANOVA indicated significant main effects of group, time, and interaction (time × group) (*P* < 0.05) in the arthrogenic contracture (°), MDA (nmol/mg), SOD (U/mg),p-Smad2/3+ cells, collagen density (%), and average protein levels.

### 3.1 Arthrogenic contracture

The arthrogenic contracture in the six groups of rabbits was shown in Table 1 and Figure 1. B. d (group C: 0°, group IM-6w: 55.33 ± 1.21°, group RM-1w: 59.50 ± 1.22°, group LIPUS-1w: 52.50 ± 1.05°, group RM-2w: 53.17 ± 0.75°, group LIPUS-2w: 46.00 ± 0.89°). As compared with group C, the arthrogenic contracture in group IM-6w was significantly greater (P < 0.05, Table 1, Figure 1. B. d). A serious arthrogenic contracture developed after 6 weeks of immobilization. With the prolongation of recovery time, both groups LIPUS and RM showed decreased arthrogenic contracture (P < 0.05, Table 1, Figure 1. B. d). A significant reduction in arthrogenic contracture was observed in the groups LIPUS compared to the groups RM at the same point in time (P < 0.05, Table 1, Figure 1. B. d).

**Table 1.** Raw values of arthrogenic contracture (°), MDA (nmol/mg), SOD (U/mg), p-Smad2/3 positive cells, collagen density (%), and average protein levels and main effect.

| Results | Group | Recovery time (week) |  |  | Main effect |  |  |  |
| --- | --- | --- | --- | --- | --- | --- | --- | --- |
|  |  | 0 | 1 | 2 |  | group | time | interaction |
| arthrogenic | LIPUS | 55.33 ± 1.21 | 52.50 ± 1.05 | 46.00 ± 0.89 | *F* | 405.899 | 72.466 | 72.280 |
| contracture (°) | RM |  | 59.50 ± 1.22 | 53.17 ± 0.75 | *P* | < 0.001 | < 0.001 | < 0.001 |
| MDA (nmol/mg) | LIPUS | 37.95 ± 0.70 | 35.23 ± 0.70 | 30.26 ± 0.47 | *F* | 1441.596 | 30.132 | 339.428 |
|  | RM |  | 43.23 ± 0.60 | 42.18 ± 1.16 | *P* | < 0.001 | < 0.001 | < 0.001 |
| SOD (U/mg) | LIPUS | 7.23 ± 0.17 | 8.13 ± 0.20 | 11.08 ± 0.14 | *F* | 1986.426 | 220.624 | 1108.747 |
|  | RM |  | 6.04 ± 0.18 | 6.06 ± 0.12 | *P* | < 0.001 | < 0.001 | < 0.001 |
| p-Smad2/3+ cells | LIPUS | 71.67 ± 0.52 | 67.00 ± 0.89 | 42.17 ± 0.75 | *F* | 23805.000 | 3551.833 | 16931.364 |
|  | RM |  | 80.67 ± 0.82 | 80.00 ± 1.10 | *P* | < 0.001 | < 0.001 | < 0.001 |
| collagen density (%) | LIPUS | 47.51 ± 0.74 | 32.11 ± 0.68 | 25.66 ± 0.90 | *F* | 17211.376 | 78.091 | 2672.453 |
|  | RM |  | 72.62 ± 0.94 | 71.64 ± 1.27 | *P* | < 0.001 | < 0.001 | < 0.001 |
| TGF-β1 / GAPDH | LIPUS | 0.56 ± 0.03 | 0.44 ± 0.03 | 0.28 ± 0.03 | *F* | 641.373 | 14.933 | 258.129 |
|  | RM |  | 0.87 ± 0.02 | 0.89 ± 0.08 | *P* | < 0.001 | 0.001 | < 0.001 |
| p-Smad2 / Smad2 | LIPUS | 0.81 ± 0.02 | 0.60 ± 0.02 | 0.49 ± 0.01 | *F* | 1906.672 | 7.861 | 132.128 |
|  | RM |  | 1.02 ± 0.05 | 1.01 ± 0.07 | *P* | < 0.001 | 0.009 | < 0.001 |
| p-Smad3 / Smad3 | LIPUS | 0.75 ± 0.01 | 0.63 ± 0.02 | 0.51 ± 0.01 | *F* | 797.101 | 71.196 | 270.264 |
|  | RM |  | 0.81 ± 0.01 | 0.82 ± 0.04 | *P* | < 0.001 | < 0.001 | < 0.001 |
| Smad4 / GAPDH | LIPUS | 0.81 ± 0.01 | 0.52 ± 0.01 | 0.39 ± 0.02 | *F* | 4542.716 | 31.477 | 425.062 |
|  | RM |  | 1.05 ± 0.06 | 1.06 ± 0.03 | *P* | < 0.001 | < 0.001 | < 0.001 |
| Collagen I / GAPDH | LIPUS | 0.87 ± 0.01 | 0.72 ± 0.02 | 0.52 ± 0.02 | *F* | 1696.602 | 124.071 | 604.542 |
|  | RM |  | 0.92 ± 0.01 | 0.91 ± 0.03 | *P* | < 0.001 | < 0.001 | < 0.001 |
| α-SMA / GAPDH | LIPUS | 0.76 ± 0.01 | 0.50 ± 0.02 | 0.34 ± 0.01 | *F* | 2552.413 | 269.165 | 1738.662 |
|  | RM |  | 0.83 ± 0.02 | 0.82 ± 0.01 | *P* | < 0.001 | < 0.001 | < 0.001 |
| Collagen III / | LIPUS | 0.78 ± 0.02 | 0.57 ± 0.02 | 0.36 ± 0.02 | *F* | 3182.987 | 173.040 | 1100.223 |
| GAPDH | RM |  | 0.93 ± 0.01 | 0.91 ± 0.01 | *P* | < 0.001 | < 0.001 | < 0.001 |

Repeated measures ANOVA indicated significant main effects of group, time, and interaction (time × group) ( $P < 0.05$ ) in the arthrogenic contracture ( $^{\circ}$ ), MDA (nmol/mg), SOD (U/mg), p-Smad2/3+ cells, collagen density (%), and average protein levels.

### 3.2. Anterior knee joint capsule fibrosis

Masson staining of the anterior knee joint capsule was performed to assess the effects of LIPUS on joint capsule fibrosis. As presented in Table 1 and Figure 2. A, high area of collagen deposition was observed in the group IM-6w by compared with group C (*P* < 0.05). In groups RM, collagen density increased significantly during remobilization compared to group IM-6w (*P* < 0.05, Table 1, Figure 2. A. b). Nevertheless, there was no significant difference between groups RM-1w and RM-2w in terms of collagen density (*P* > 0.05, Table 1, Figure 2. A. b). It was also found that with the prolongation of treatment time, collagen density in groups LIPUS showed a clearly reducing trend. Compared to group LIPUS-1w, group LIPUS-2w showed a more obvious reduction (*P* < 0.05, Table 1, Figure 2. A. b).

**Figure 2.**
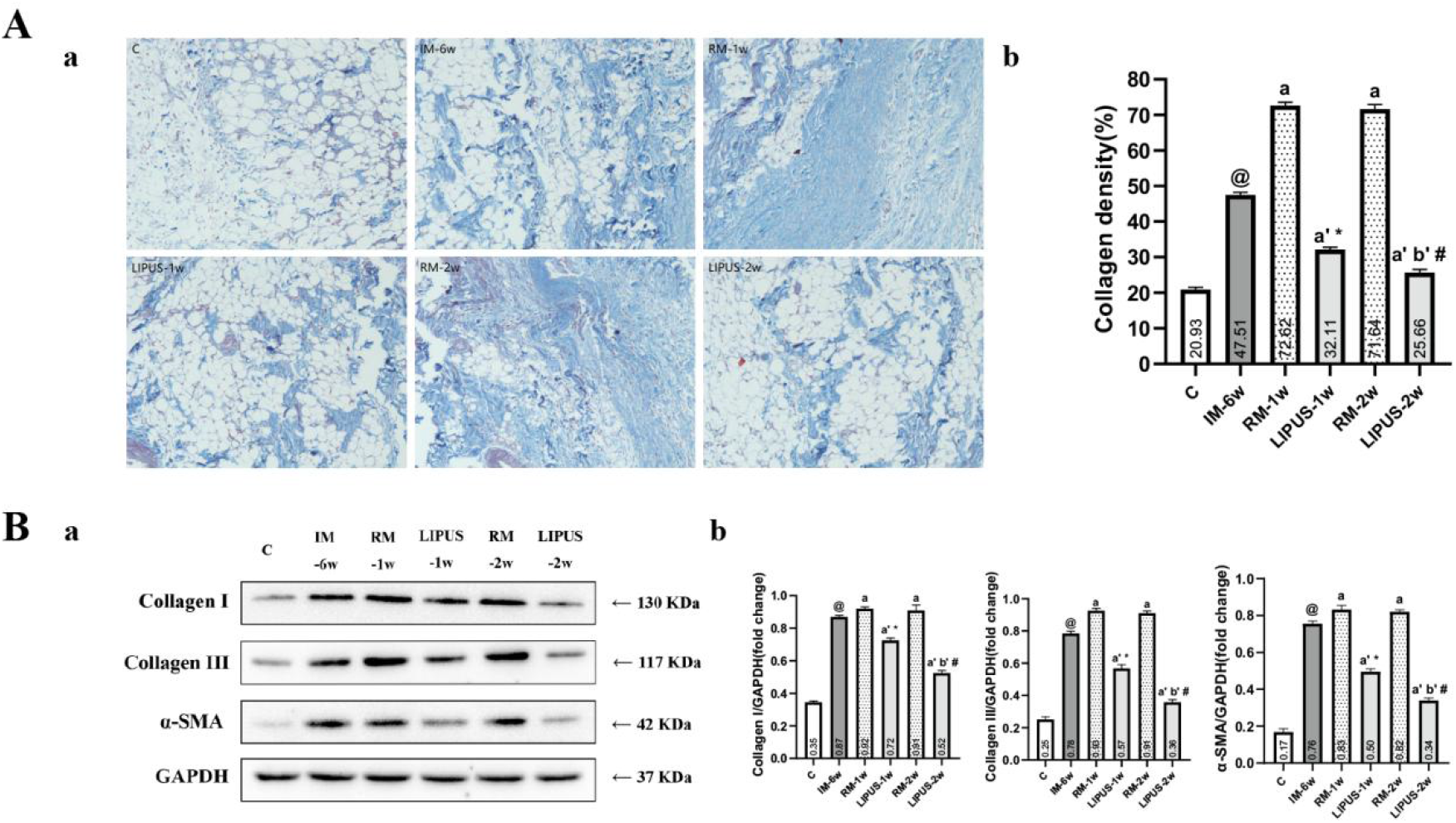
(A) (a)Collagen relative area of joint capsule in each group (n=6). (b) The effects of remobilization and LIPUS on the relative area of the collagen. ^@^ *P* < .05 vs. group C, ^a^ *P* < .05 vs. group IM-6 w, ^a’^ *P* < .05 vs. group IM-6 w, ^b’^ *P* < 0.05 vs. group LIPUS-1 w, ^*^ *P* < .05 vs. RM-1 w, ^#^ *P* < .05 vs. group RM-2 w.(B) (a) The expression of collagen I, collagen III and α-SMA in the anterior joint capsule were examined by Western blotting in each group (n=6). (b)The average protein level of Collagen I/GAPDH, Collagen III/ GAPDH and α-SMA/GAPDH of the joint capsule. ^@^ *P* < .05 vs. group C, ^a^ *P* < .05 vs. group IM-6w, ^b^ *P* < .05 vs. group RM-1w, ^a’^ *P* < .05 vs. group IM-6w, ^b’^ *P* < .05 vs. group LIPUS-1w, ^*^ *P* < .05 vs. group RM-1w, ^#^ *P* < .05 vs. Group RM-2w.

Then, we investigated the expression of Collagen I, Collagen III and α-SMA in anterior knee joint capsule to further confirm the results of joint capsule fibrosis. As presented in Table 1 and Figure 2.B, the average protein levels for Collagen I, Collagen III and α-SMA were significantly greater in group IM-6w compared with group C (*P* < 0.05, Table 1, Figure 2. B. b). Similar to the results of collagen density, the average protein levels for Collagen I, Collagen III and α-SMA were significantly higher in groups RM than in group IM-6w (*P* < 0.05, Table 1, Figure 2. B. b), However, there was no significant difference between groups RM-1w and RM-2w (*P* > 0.05, Table 1, Figure 2. B. b). Importantly, in the results of groups LIPUS, an obviously reducing trend of the average protein levels for Collagen I, Collagen III and α-SMA was observed with the prolongation of treatment time, and the reduction in group LIPUS-2w was more obvious than that in group LIPUS-1w (P < 0.05, Table 1, Figure 2. B. b).

### 3.3 The level of ROS

By collecting the supernatants from the anterior knee joint capsule at the previously mentioned time points, changes in MDA and SOD levels were assessed after LIPUS application. Compared with the group C, SOD of the group IM-6w significantly decreased while MDA increased significantly (*P* < 0.05, Table 1, Figure 3. A). In the groups RM, SOD levels significantly decreased and MDA levels significantly increased compared with the group IM-6w(*P* < 0.05, Table 1, Figure 3. A). Nonetheless, there was no significant difference in the levels of MDA and SOD between group RM-1w and RM-2w (*P* > 0.05, Table 1, Figure 3. A). SOD levels increases with treatment duration, MDA levels decreases with treatment duration. As shown in Figure 3. A, SOD levels in group LIPUS-1w were lower than those in group LIPUS-2w, while the level of MDA in group LIPUS-1w was higher than that in group LIPUS-2w (*P* < 0.05, Table 1). In the same time point, SOD in groups RM was lower than that in groups LIPUS, whereas MDA in groups RM was higher (*P* > 0.05, Table 1, Figure 3. A).

**Figure 3.**
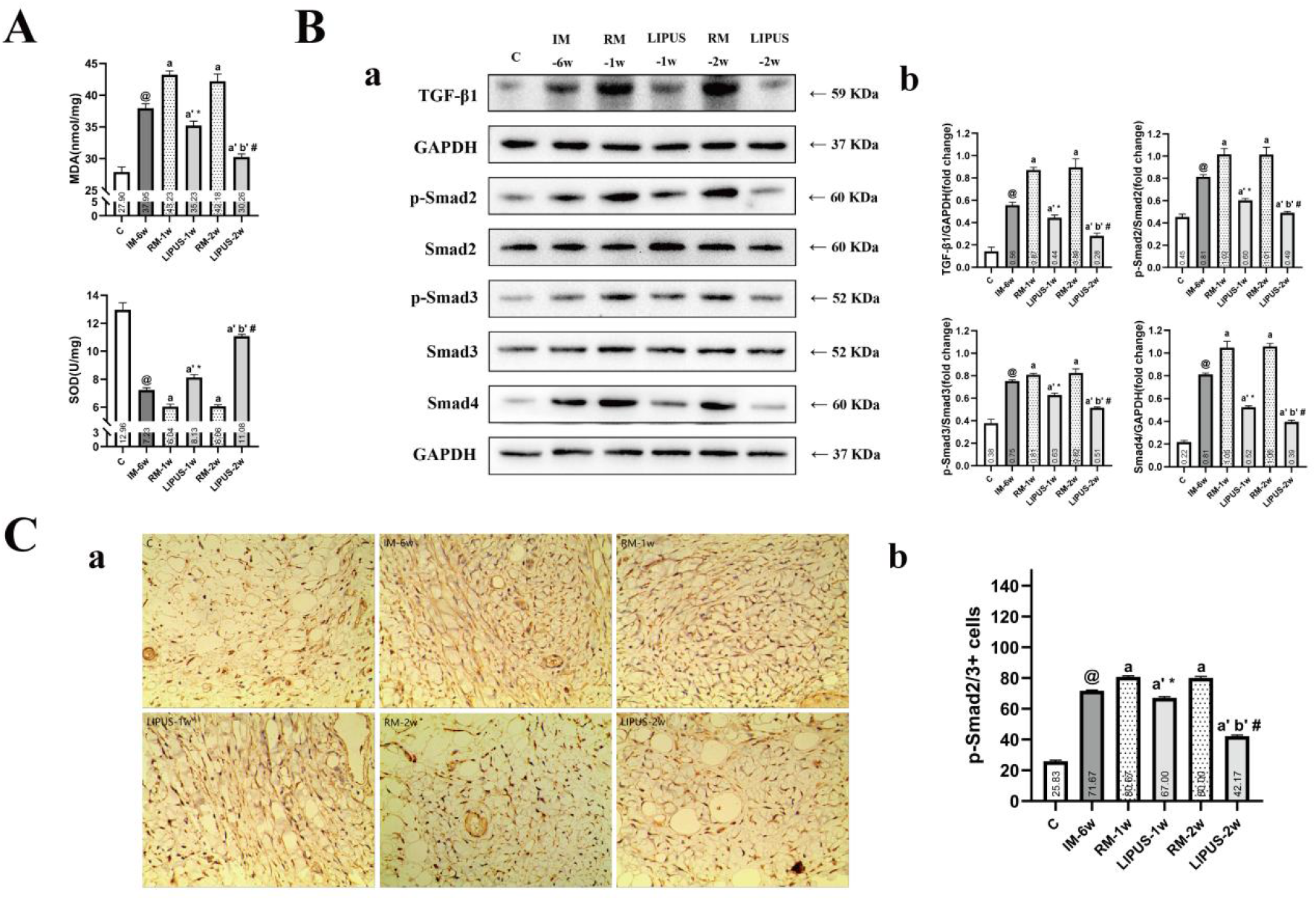
(A) Effects of LIPUS on the levels of SOD and MDA in each group (n=6). ^@^ *P* < .05 vs. group C, ^a^ *P* < .05 vs. group IM-6w, ^b^ *P* < .05 vs. group RM-1w, ^a’^ *P* < .05 vs. group IM-6w, ^b’^ *P* < .05 vs. group LIPUS-1w, ^*^ *P* < .05 vs. group RM-1w, ^#^ *P* < .05 vs. group RM-2w. (B) (a) Expression of TGF-β1/Smad signaling pathway in the anterior joint capsule were examined by Western blotting in each group (n=6). (b) The average protein level of TGF-β1/GAPDH, p-Smad2/Smad2, p-Smad3/Smad3, smad4/GAPDH, in the anterior joint capsule. ^@^ *P* < .05 vs. group C, ^a^ *P* < .05 vs. group IM-6w, ^b^ *P* < .05 vs. group RM-1w, ^a’^ *P* < .05 vs. group IM-6w, ^b’^ *P* < .05 vs. group LIPUS-1w, ^*^ *P* < .05 vs. group RM-1w, ^#^ *P* < .05 vs. group RM-2w. (C) (a) Immunohistochemical staining for p-Smad2/3 positive cells of the anterior joint capsule in each group (n=6). (b) Quantification of p-Smad2/3 positive cells in the anterior joint capsule. ^@^ *P* < .05 vs. group C, ^a^ *P* < .05 vs. group IM-6w, ^b^ *P* < .05 vs. group RM-1w, ^a’^ *P* < .05 vs. group IM-6w, ^b’^ *P* < .05 vs. group LIPUS-1w, ^*^ *P* < .05 vs. group RM-1w, ^#^ *P* < .05 vs. group RM-2w.

### 3.4 The activation of TGF-β1/Smad signaling pathway

The activation of the TGF-1 signaling cascade is known to contribute to the initiation and development of fibrosis, particularly Smad2, Smad3 and Smad4. In this study, we analyzed the effects of LIPUS and remobilization on the expression of the TGF-1 signaling pathway protein in the anterior knee joint capsule. The expression of TGF-β1, Smad2, p-Smad2, Smad3, p-Smad3 and Smad4 in anterior knee joint capsule were detected by western blot (Table 1, Figure 3. B. a). According to western blot analysis, the expression of TGF-β1, p-Smad2/ Smad2, p-Smad3/Smad3 and Smad4 in group IM-6w was upregulated by compared with group C, indicating that TGF-β1/Smad signaling pathway was activated. (*P* < 0.05, Table 1, Figure 3. B. b). Although the average protein levels for these also increased significantly in groups RM compared with group IM-6w, but there were no significant differences between groups RM-1w and RM-2w. (*P* > 0.05, Table 1, Figure 3. B. b). Importantly, TGF-β1/Smad signaling pathway protein was observed more obvious effect of inhibition in groups LIPUS with the prolongation of treatment time. When groups LIPUS and RM were compared at the same timepoint, TGF-1/Smad signaling pathway protein expression was significantly lower in LIPUS than in RM (*P* < 0.05, Table 1, Figure 3. B).

Furthermore, p-Smad2/3 positive cells were also indicative of an activation of the TGF-β1/Smad signaling pathway. As shown in Table 1 and Figure 3.C, we found that the number of p-Smad2/3 positive cells was obviously increased in group IM-6w by compared with group C (*P* < 0.05, Table 1, Figure 3. C. b). Furthermore, obtained results also revealed considerably high number of p-Smad2/3 positive cells in groups RM, whereas little p-Smad2/3 expression was observed in groups LIPUS (*P* < 0.05, Table 1, Figure 3. C. b). Although there was no difference in number of cells between group RM-1w and group RM-2w (*P* > 0.05, Table 1, Figure 3. C. b), the cells of group LIPUS-2w was less than that of LIPUS-1w (*P* < 0.05, Table 1, Figure 3. C. b). Thus, LIPUS efficiently inhibited the TGF-β1/Smad signaling pathway in anterior knee joint capsule.

## 4. Discussion

With the development of joint contracture research, the role of joint capsule fibrosis in the progression of joint contracture has been widely concerned (Zhou et al. 2020). There is a general view that arthrogenic contracture mainly induced by joint capsule fibrosis is aroused by long-term immobilization (Zhou et al. 2020; Zhang et al. 2018). Clinically, it is widely accepted that facilitation of joint movements improves immobilization-induced joint contractures. Previous research, however, indicated that immobility-induced restriction of joint motion was not fully restored after remobilization (Ando et al. 2012). Apart from this, previous research results suggested that active exercises during an early period of remobilization even partly hinder recovery from contractures caused by immobilization (Kaneguchi et al. 2018; Kaneguchi et al. 2018). As a result, it is necessary to find an effective therapy to reduce joint capsule fibrosis during remobilization. Recently, the role of LIPUS in anti-fibrosis has received great attention (Aibara et al. 2020; Liao et al. 2021; Zhao et al. 2021). Herein, we demonstrated that LIPUS can suppress anterior knee joint capsule fibrosis and arthrogenic contracture through inhibiting TGF-β1/Smad signaling pathway during remmobilization, thus indicating the potential of LIPUS as a novel therapeutic method for joint contracture.

In agreement with our previous studies (Zhou et al. 2020), the present study showed that arthrogenic contracture in group IM-6w were greater than those in groups C. It was evident from Masson staining that collagen deposition increased after 6 weeks of immobilization, indicating severe anterior knee capsule fibrosis. Correspondingly, our data also showed that the protein expression of Collagen I, Collagen III and α-SMA were significantly increased after 6 wks-immobilization. Activated fibroblasts and myofibroblasts synthesize ECM proteins such as collagen and α-SMA (Tokuda et al. 2021). Previous studies have shown that TGF-β1 is the major profibrotic cytokine in the fibrosis process, and the increased activity of TGF-β1 was associated with joint capsule fibrosis, whereas various pharmacological inhibitor of TGF-β receptor I kinase gave remarkable protective effects in knee joint contracture models (Mao et al. 2021; Zhou et al. 2020). TGF-β1 signaling of myofibroblast differentiation is known to occur through two pathways: Smad and ERK (Attisano et al. 2002; Hough et al. 2012). Details of these transduction pathways have proved to be complex, may vary in different cell types, and remain incompletely understood (Barnes et al. 2011; Hinz et al. 2007). In the present study, a significant amount of TGF-β1 was detected in anterior knee joint capsule preparation after immobilization. Consistently, the protein expression of p-Smad2/Smad2, p-Smad3/Smad3 and Smad4 was also upregulated in group IM-6w. In addition, as present in Figure, increased phosphorylation level of Smad2/3 manifested as an increase in the number of phosphorylated Smad2/3 positive cells. These results and those of previous study collectively indicate that TGF-1 induces joint capsule fibrosis as assessed by α-SMA expression and collagen deposition via a signaling cascade involving the TGF-1 receptor (TGF-1R)1/Smad pathway (Mao et al. 2021; Tokuda et al. 2021; Xiao et al. 2020).

Myogenic and arthrogenic contractures were classified as types of joint contractures. According to our previous study, remobilization after 4 weeks-immobilization partially recovered overall ROM by improving myogenic contracture (Wang et al. 2022). However, whether remobilization drives the worsening of arthrogenic contracture is still controversial. Previous research indicated that remobilization after 3 wks-immobilization aggravated arthrogenic contracture through inflammation and subsequent fibrosis in the joint capsule (Ando et al. 2012). Another study also showed that remobilization following short-term (1 or 2 weeks) immobilization improved myogenic contracture, but developed arthrogenic contracture (Trudel et al. 2014). Our study indicated that 1 week of immobilization instead of 2 weeks of immobilization aggravated arthrogenic contracture severity. Connective tissue stiffness, which may affect the extensibility of the joint capsule and skeletal muscle, is determined by various interacting factors, including collagen content and cross-sectional area (Wang et al. 2022; Trudel et al. 2014; Zhou et al. 2020). Type I and III collagen protein expressions and collagen density were significantly higher in group RM-1w than in group IM-6w after 1 week of remobilization, however, there was no significant difference between group RM-1w and group RM-2w in those. In previous research, researchers found that other joint components, such as ligaments, are also affected by joint immobilization and remobilization, potentially contributing to arthrogenic contracture (Trudel et al. 2014). As a result, we assumed the reduction in arthrogenic contracture in group RM-2w was due to this. Furthermore, a previous study reported that ROM limitation of rat knee joint induced by 2 wks-hindlimb immobilization was decreased by 7 wks-active exercise (Morimoto et al. 2013). In summary, we speculated both the duration of immobilization and remobilization may also influence arthrogenic contracture recovery.

In conclusion, LIPUS would be a very useful biophysical stimulus for different musculoskeletal disorders. LIPUS is a type of mechanical energy carried by sound waves over the hearing limits of the human ear. The LIPUS produces vibration forces in all tissue components, both intracellular and extracellular, delivering energy as acoustic pressure waves. Numerous studies have examined the effects of LIPUS on fibrosis. Previous study reported that daily LIPUS therapy ameliorated inflammation and tubulointerstitial fibrosis in experimental hypertensive nephropathy and diabetic nephropathy by directly inhibited the TGF-β1/Smad signaling pathway (Aibara et al. 2020). Apart from this, previous research also found that LIPUS inhibited fibroblast-like synoviocyte proliferation via downregulating Wnt/β-catenin pathway to manage synovial fibrosis in osteoarthritis (Liao et al. 2021). However, few studies have investigated the effect of LIPUS on joint capsule fibrosis induced by immobilization. Importantly, LIPUS plays a very important role in suppressing ROS release by several cells, such as neurons, endothelial cell and cartilage cells (Chen et al. 2021; Li et al. 2018; Zuo et al. 2021). Previous research demonstrated that high ROS generation not only elevated the expression of TGF-β1 but also increased the fibrosis protein markers including collagen I and α-SMA in synovial myofibroblasts after knee joint immobilization (Siani et al. 2014). In our present study, LIPUS significantly repressed the severity of anterior knee joint capsule fibrosis as evidenced by reduced arthrogenic contracture and collagen content. Furthermore, LIPUS increased SOD activity, and thus reduced the MDA content in anterior knee joint capsule, which associated with decreased formation of ROS. As expected, LIPUS lowered the expression of TGF-β1, the phosphorylation level of Smad2/3 and the expression of Smad4 in anterior knee joint capsule. Besides, the results of immunohistochemical staining for p-Smad2/3 positive cells of the anterior joint capsule provided further evidence for the inhibitory effect for TGF-β1/Smad signaling. Collagen I, Collagen III and α-SMA were the major components of the connective tissue in fibrotic septa, which is mainly produced by myofibroblasts (Wang et al. 2022). Consistently, the upregulation of Collagen I, Collagen III and a-SMA was also suppressed in groups LIPUS.

To our knowledge, this is the first study that demonstrated LIPUS attenuated the severity of immobilization-induced arthrogenic contracture and capsule fibrosis, but this study also has certain limitations. Joint capsule extensibility is generally determined by length and stiffness. Due to the limitation of experimental conditions, we did not assess the degree of length and stiffness of joint capsule. According to the method described by Zhou (2018) et al, AFMG-stained sections should be performed to measure the synovial lengths of the joint capsule. In addition, immobilization and remobilization-induced ROS generation was not directly quantified. DCFH-DA staining and flow cytometry could display a more intuitive visual representation of the ROS level. At last, precision therapy is thought to be the direction of future treatment strategies, and future studies are needed to better define optimal treatment localization.

## 5. Conclusions

In conclusion, we demonstrated that high level ROS and hyperactive TGF-β1/Smad signaling pathway might be one of the causes of knee joint capsule fibrosis induced by immobilization and LIPUS attenuated the severity of immobilization-induced knee joint capsule fibrosis through inhibition of the production of ROS and the activation of TGF-β1/Smad signaling pathway. Based on our experimental findings, we hypothesize that LIPUS could be a promising therapeutic modality to ameliorate arthrogenic contracture and joint capsule fibrosis after long-term immobilization in human.

## Acknowledgments

The authors thank Professor *Hua Wang* from the School of Public Health, Anhui Medical University for his valuable guidance and corrections.

